# Toward an Over-parameterized Direct-Fit Model of Visual Perception

**DOI:** 10.1101/2022.10.06.511127

**Authors:** Xin Li

## Abstract

In this paper, we revisit the problem of computational modeling of simple and complex cells for an over-parameterized and direct-fit model of visual perception. Unlike conventional wisdom, we highlight the difference in parallel and sequential binding mechanisms between simple and complex cells. A new proposal for abstracting them into space partitioning and composition is developed as the foundation of our new hierarchical construction. Our construction can be interpreted as a product topology-based generalization of the existing k-d tree, making it suitable for brute-force direct-fit in a high-dimensional space. The constructed model has been applied to several classical experiments in neuroscience and psychology. We provide an anti-sparse coding interpretation of the constructed vision model and show how it leads to a dynamic programming (DP)-like approximate nearest-neighbor search based on 𝓁_∞_-optimization. We also briefly discuss two possible implementations based on asymmetrical (decoder matters more) auto-encoder and spiking neural networks (SNN), respectively.

## I. Introduction

How do we learn to see in the first six months after birth? To answer this question, David Hubel and Torsten Wiesel conducted pioneering experiments in the 1950s, leading to the discovery of simple and complex cells [43]. Inspired by their discovery, David Marr developed a theory of the neocortex [62] in 1970 and a theory of the hippocampus [63] in 1971. His computational investigation of vision [61] was published in 1982 after his death. The construction of neocognitron by Fukushima [31] and connectionist models by LeCun [50] in the 1980s represented the continuing effort to construct biologically plausible computational models for visual perception. Wavelet theory [57, 23] and sparse coding [70, 71] in the 1990s further supplied mathematical formulations of multi-resolution analysis for scale-invariant representation of images. Rapid advances in deep learning [33, 51], especially the class of over-parameterized models [7, 6] have expedited both the theory and practice of data-driven/learning-based visual processing.

Despite the great progress of today, the gap between biological and artificial vision remains significant in the following aspects. First, the network architecture of the convolutional neural network (CNN) is characterized by the pooling of layers, which reduces the dimensionality of the input data. This is in sharp contrast to the increase in the number of neurons and synapses as we move from the lower layer (e.g. V1) to the higher layer (e.g. V4) of neocortex. This anatomical finding has inspired H. Barlow to revise his redundancy reduction hypothesis into the redundancy exploitation hypothesis [9] in 2001. Second, although recurrent neural networks including long short-term memory (LSTM) [42] take into account temporal dynamics and have found successful applications in 1D signal analysis, the role of memory in visual perception has remained explored. In the human vision system, the hippocampus is known to play a critical role in various cognition tasks, including memory consolidation and novelty detection [49]. Finally, it remains a mystery how the human brain can manage to achieve the objectives of learning and memory with more than 100-1000 trillion synapses and a power budget of less than 20W. The challenge of breaking the conventional von Neumann architecture built upon the Turing machine remains a holy grail in neuromorphic computing.

The motivation behind this paper is two-fold. On the one hand, both the human brain and CNN are characterized by the ability to optimize an astronomical number of synaptic weights [20]. The class of over-parameterized models [7, 6] has shown some counterintuitive properties, such as double descent [68]. Analytical tools such as neural tangent kernel (NTK) offer an approach to understanding over-parameterization in the Hilbert space, but, like all kernel methods, they are not compatible with the recursion strategy (e.g. dynamic programming that builds upon the optimality of substructures [13]). We seek to understand overparameterization in the framework of optimizing hierarchical representations [85]. On the other hand, an evolutionary perspective on biological and artificial neural networks [34] offers a direct-fit approach in brute force. Such a deceivingly simple model, when combined with over-parameterized optimization, offers an appealing solution to increase the generalization (predictive) power without explicitly modeling the unknown generative structure underlying sensory inputs.

In this paper, we construct an over-parameterized direct-fit model for visual perception. Unlike the conventional wisdom of abstracting simple and complex cells, we use space partitioning and composition as the building block of our hierarchical construction. In addition to biological plausibility, we offer a geometric analysis of our construction in topological space (i.e., topological manifolds without the definition of a distance metric or an inner product). Our construction can be interpreted as a product-topology-based generalization of the existing k-d tree [14], making it suitable for brute-force direct-fit in a high-dimensional space. In the presence of novelty/anomaly, a surrogate model that mimics the escape mechanism of the hippocampus can be activated for unsupervised continual learning [98]. The constructed model has been applied to several classical experiments in neuroscience and psychology. We also provide an anti-sparse coding interpretation [46] of the constructed vision model and present a dynamic programming (DP)-like solution to approximate nearest neighbor in high-dimensional space. Finally, we briefly discuss two possible network implementations of the proposed model based on asymmetric autoencoder [69] and spiking neural networks (SNN) [45], respectively.

## II. Neuroscience Foundation

### A. Dichotomy: Excitatory and Inhibitory Neurons

During the work of Wilson and Cowan in the 1970s [92, 93], they made the crucial assumption that “all nervous processes of any complexity depend on the interaction of excitatory and inhibitory cells.” Using phase plane methods, they have shown simple and multiple hysteresis phenomena and limit cycle activity with localized populations of model neurons. Their results, more or less, offer the primitive basis for memory storage, namely stimulus intensity, which can be coded in both the average spike frequency and the frequency of periodic variations in the average spike frequency [78]. However, such ad hoc sensory encoding cannot explain the sophistication of learning, memory, and recognition associated with higher functions.

### B. Hebbian Learning and Anti-Hebbian Learning

Hebbian learning [38] is a dogma that claims that an increase in synaptic efficacy arises from repeated and persistent stimulation of a presynaptic cell by a postsynaptic cell. Hebbian learning rule is often summarized as “cells that fire together wire together”. The physical implementation of the Hebbian learning rule has been well studied in the literature, for example, through spike timing-dependent plasticity (STDP) [16]. The mechanism of STDP is to adjust the connection strengths based on the relative timing of some neuron’s input and output action potentials. STDP as a Hebbian synaptic learning rule has been demonstrated in various neural circuits, from insects to humans.

By analogy to excitatory and inhibitory neurons, it has been suggested that a reversal of Hebb’s postulate, named anti-Hebbian learning, dictates the reduction (rather than increase) of the synaptic connectivity strength between neurons following a firing scenario. Synaptic plasticity that operates under the control of an anti-Hebbian learning rule has been found to occur in the cerebellum [12]. More importantly, local anti-Hebbian learning has been shown to be the foundation for forming sparse representations [27]. By connecting a layer of simple Hebbian units with modifiable anti-Hebbian feedback connections, one can learn to encode a set of patterns into a sparse representation in which statistical dependency between the elements is reduced while preserving the information. However, the sparse coding represents only a local approximation of the sensory processing machinery. To extend it to global (nonlocal) integration, we have to assume an additional postulate, called the “hierarchical organization principle “, which we will introduce in the next section.

### C. Simple and Complex Cells in V1

These two classes of cells were discovered by Torsten Wiesel and David Hubel in the early 1960s [43]. Simple cells respond primarily to oriented edges and gratings, which can be mathematically characterized by Gabor filters [24]. Complex cells also respond to oriented structures; unlike simple cells, they have a degree of spatial invariance. The difference between receptive fields and the characteristics of simple and complex cells has inspired the invention of the neocognitron by Fukushima in 1979 [31], which foresaw the subsequent convolutional neural network. The hierarchical convergent nature of visual processing has also inspired the construction of the HMAX model in 1999 [81].

An important observation with the difference between simple and complex cells, as described in [43], is their neural circuits and the corresponding temporal dynamics. Simple cells are built from center-surrounding cells, which require *simultaneous summation*. On the contrary, activation of the complex cell by a moving stimulus requires *successive activation* of many simple cells. Therefore, the spatial invariance of complex cells is achieved by summation and integration of the receptive fields of simple cells. Mathematical modeling of complex cells has been extensively studied in the literature (e.g., energy model [3]). However, the abstraction strategies taken in this paper will be different from those in the open literature.

### D. Mountcastle’s Universal Principle

In 1978 Mountcastle suggested a universal processing principle that has been acclaimed as the Rosetta Stone of neuroscience. According to Mountcastle, all parts of the neocortex operate according to a common principle, the cortical column being the unit of computation [67]. If Mountcastle were correct, the “simple discovery” made by Hubel and Wiesel might have deeper implications in the mechanism of visual processing beyond V1. Along this line of reasoning, the striking difference between simultaneous and successive activation of simple and complex cells might illustrate a fundamental contrast between two classes of binding mechanism among neurons.

In visual perception, it has been hypothesized that the characteristics of individual objects are bound / segregated by Hebbian / anti-Hebbian learning of different groups of neurons [66]. We conjecture that there exist two types of binding mechanism (parallel vs. sequential) that are analogous to combinatorial and sequential logic in digital circuits. The former plays the role of integrating spatially overlapped parts into a whole (e.g., a horizontal edge and a vertical edge form a letter “T”) or multiple features of the same object into a coherent perception (e.g., the age and gender of a face) in object recognition. Parallel binding can be interpreted as an extension of von der Malsburg’s correlation theory [58]. This is the mechanism adopted by the dorsal stream to support the task of object vision. The latter is at the core of integrating spatially non-overlapped parts into a whole (e.g., the concatenation of letters into a word) in spatial vision, which belongs to the ventral stream/pathway. Sequential binding is closely related to the formation of short-term memory (e.g., Miller’s law [65]) and long-term memory (e.g., Atkinson–Shiffrin model [8]) in the brain. The fundamental difference between parallel and sequential binding is that the former is invariant to permutation (the ordering of parts does not affect the perception of the whole), while the latter is sensitive to the ordering of the neuronal groups.

Binding by neuronal synchrony has been widely recognized in the literature; however, the binding problem is often thought to suffer from the so-called “superposition catastrophe” [91]. The combination coding argument often faces the dilemma of the curse of dimensionality, and it has been suggested that a representation with hierarchical structure can at least partially overcome this barrier of impracticality with combination coding. More importantly, we argue that our intuition about the capacity of the cortex might be misleading and our understanding of the power of hierarchical structures is inadequate [35]. If the curse of dimensionality can become a blessing [18], the combination coding can be made compatible with the hypothesis of redundancy exploitation [9].

## III. Construction of an Over-parameterized Direct-Fit Model

In neuroscience, the principle of hierarchical organization can be roughly stated as follows: The nested structure of the physical world is mirrored by the hierarchical organization of the neocortex [35]. Unlike the mathematical construction of wavelets [23, 57], we envision that nature has discovered an elegant “elementary” solution in topological space (without the extra structure of distance metric or inner product) to manage the complexity of sensory stimuli in the physical world. We propose to study the following problem as the fruit-fly problem in visual perception [39].

### Problem Formulation of Visual Perception

Given a visual stimulus (e.g., a sequence of images) as input, group/cluster them into different classes in an unsupervised manner.

The solution, as manifested by an infant’s development of the visual cortex (primarily for ventral stream for object vision) during the first six months after birth, lies in a novel construction of a hierarchical direct-fit model based on simple and complex cells. As argued by Jean Piaget [76], the ordering of mathematical spaces in early children cognitive development (topology before geometry) is the opposite to that of what we have learned in school (topology after geometry). Therefore, we attempt to construct our visual perception model in topological space with the least amount of assumed mathematical structures.

#### A. Preliminary on Topological Space

To facilitate our abstraction of simple and complex cells by subspace and product topology, we briefly review the basics of topological space as follows. We will follow the axiomatization of Felix Hausdorff to construct the topological space using neighborhood as the building block. Let 𝒩 denote the neighborhood function assigning to each point *x* ∈ **X** a non-empty subset 𝒩 (*x*) ∈ **X**. Then the following axioms must be satisfied for **X** with 𝒩 to be called a topological space.

1. If **N** is a neighborhood of *x* (i.e., **N** ∈ 𝒩 (*x*)), then *x* ∈ **N**;
2. If **N** is a subset of **X** and includes a neighborhood of *x*, then **N** is a neighborhood of *x*;
3. The intersection of two neighborhoods of *x* is a neighborhood of *x*;
4. Any neighborhood **N** of *x* includes a neighborhood **M** of *x* such that **N** is a neighborhood of each point in **M**.

Note that the fourth axiom plays the role of linking the neighborhoods of different points in **X** together. Since no distance metric is defined, we need to define the basis as the starting point for defining a topology.

##### Basis of a Topology

Let **X** be a set, and suppose that ℬ is a collection of subsets of **X**. Then ℬ is a basis for some topology in **X** if and only if the following two conditions are satisfied: a) ∪_**B**_∈B **B** = **X**; b) If **B**_1_, **B**_2_ ∈ ℬ and *x* ∈ **B**_1_ ∩ **B**_2_, there exists an element **B**_3_ ∈ ℬ such that *x* ∈ **B**_3_ ⊆ **B**_1_ ∩ **B**_2_.

In this work, we will only consider the neighborhood basis, which is defined by

##### Neighborhood Basis

A neighborhood basis is a subset ℬ ⊆ 𝒩 (*x*) such that for all **V** ∈ 𝒩 (*x*), there exists some *B* ∈ ℬ such that **B** ⊆ **V**. In other words, for any neighborhood **V** we can find a neighborhood **B** on the basis of the neighborhood contained in **V**.

With the above setup, the objective is to construct a hierarchical direct-fit model in the Hausdorff space (a.k.a. topological manifold), which generalizes the existing multi-resolution analysis in the Hilbert space [57]. Following our intuition above, simple and complex cells will be abstracted into subspace and product topology [54], respectively. Formally, we have the following.

##### Subspace Topology

Let **X** be a topological space and let **S** ⊆ **X** be any subset. Then 𝒯_*S*_ = {**U** ⊆ **S** : **U** = **S** ∩ **V** *for some open subset* **V** ⊆ **X**} is the subspace topology.

##### Product Topology

Suppose that **X**_1_, …, **X**_*n*_ are arbitrary topological spaces. In its Cartesian product **X**_1_ × … × **X**_*n*_, the product topology is generated on the following basis: ℬ = {**U**_1_ × … ×**U**_*n*_ : **U**_*i*_ *is an open subset of* **X**_*i*_, *i* = 1, …, *n*}.

Both subspace and product topologies have their uniqueness in terms of satisfying the characteristic property [48]. The geometric intuition behind our construction of the new hierarchical model is best illustrated by the duality between space partitioning (i.e., subspace topology) and composition (i.e., product topology).

#### B. Computational Modeling of Neocortex

**Simple Cells** play the role of space partitioning, which can be abstracted as subspace topology [48]. A good proxy model to study the concept of space partitioning is the k-dimensional tree (k-d tree) [14], a space partitioning data structure to organize points in a k-dimensional space. The k-d tree structure can be interpreted as a class of binary space partitioning trees that extends the binary search tree (BST) for sorted arrays. It directly fits the data using hyperplanes to recursively partition the k-dimensional space. A simple variant of the k-d tree, named the random projection tree (rp tree) [22], is capable of automatically adapting to the low-dimensional structure of the data without explicit manifold learning.

The limitations of linear separability (e.g., single-layer perceptron [82]) are well known. Conventional wisdom of addressing these limitations is to introduce a nonlinear activation unit or a hidden layer (e.g., multi-layer perceptron). However, we note that some linearly non-separable set (e.g., the well-known XOR as shown in Fig. 1) can be decomposed into the linear difference between two linearly separable sets (*X* ⊕ *Y* = (*X*|*Y*) − (*X* ∧ *Y*)). Such an observation, when combined with the data structure such as k-d/rp trees, offers a refreshing perspective of adapting to the intrinsic low-dimensional structure in high-dimensional data. That is, instead of nonlinear dimensionality reduction (e.g., the well-known Johnson-Lindenstrauss lemma [89]), we can use only a linear combination of space partitioning and differencing to approximate an arbitrary nonconvex object based on the following lemma.

**Fig. 1:**
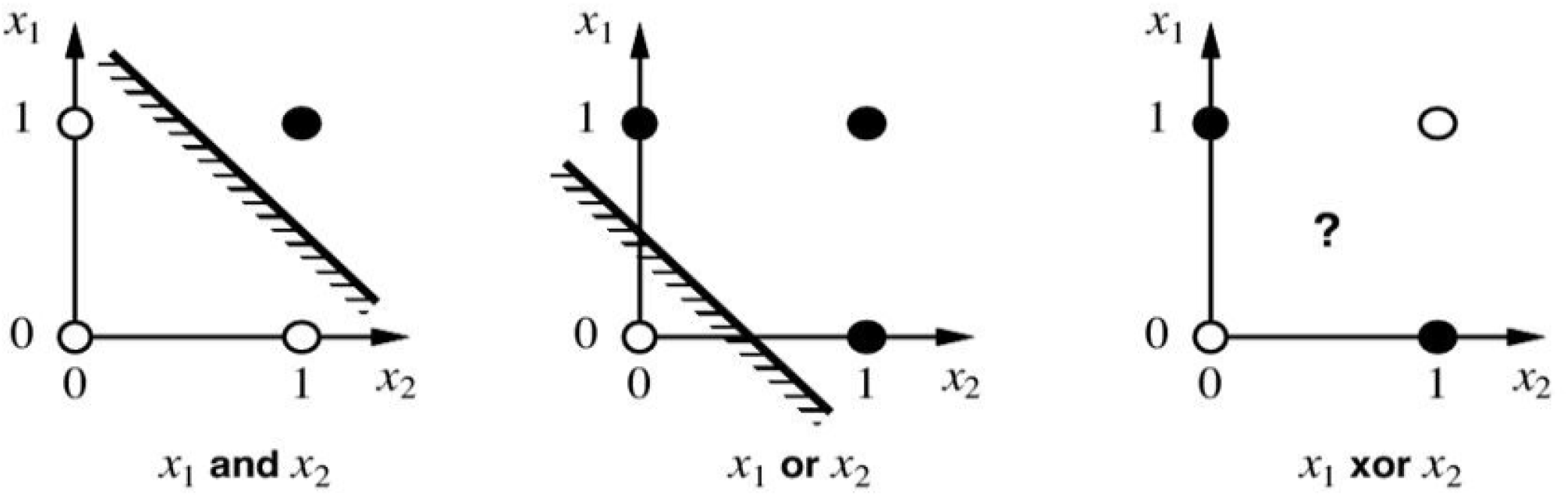
XOR is not linearly separable but can be decomposed into the linear difference between linearly separable sets (the base case for proving the convex decomposition lemma).

##### Lemma 1

Convex Decomposition Lemma for Simple Cells

Any nonconvex object *X* can be decomposed into a finite set of differences between convex objects. **Sketch of the Proof** Prove by induction with *n*, which is the minimum number of convex objects {*X*_1_, …, *X*_*n*_} for the generation of *X*. Starting with *n* = 2, it is easy to construct the solution for *X* = *X*_1_±*X*_2_. Suppose that the statement is true for the case of *n* = *k*, then the solution to *n* = *k* + 1 can be constructed similarly to *n* = 2.

Note that k-d trees have the nice property of facilitating NN/kNN search in metric space [14, 29]; but its performance degrades in high-dimensional space due to the curse of dimensionality. More importantly, one of the surprising behavior of distance metrics in high-dimensional spaces [4] is that 𝓁_*p*_ (*p* → 0) behaves much better than the popular Euclidean distance metric. Counterintuitively, in a high-dimensional space, proximity-based queries such as the NN search are meaningless and unstable because the discrimination between the nearest and farthest neighbor becomes poor. As rigorously shown in [4], the relative contrast provided by a norm with a smaller parameter *p* is more likely to dominate another norm with a larger parameter *p* as the dimensionality increases. The fractional distance concentration [28] dictates that we either work with topological space (instead of metric space) or use 𝓁_0_ as a pseudo-norm that recognizes the limitations of distance metrics. Interestingly, the problem of 𝓁_∞_-optimization has shown to be useful for an approximate NN search with anti-sparse coding [46].

**Complex Cells** play the role of the space composition, which can be abstracted by product topology [48]. Note that our intuition is consistent with the max/sum-pooling operation in HMAX model [81] because the objective is to achieve spatial invariance within an increased field of view. The difference lies in the way of abstraction - from simple to complex cells, we ask what will be its dual operation of space partitioning. Along this line of reasoning, if simple cells are responsible for linear separability without the change of dimensionality; complex cell must be able to increase the dimensionality for handling more sophisticated objects.

A common criticism of increasing dimensionality is concern about the so-called *curse of dimensionality* [13]. There are several ways to address this problem. First, recently proposed overparameterized models [7] or direct-fit models [34] suggest that dimensionality can be a blessing when a large amount of training data is available due to a counterintuitive phenomenon called “concentration of measure” [52]. Second, experimental studies such as [18] have demonstrated the blessing of dimensionality in face verification applications, which is consistent with biological findings [64]. More importantly, the low-dimensional manifold structure can still be preserved after space composition because of the following lemma.

##### Lemma 2

Product-Manifold Lemma [54] for Complex Cells

In topology, a topological manifold is a topological space that locally resembles the real *n*-dimensional Euclidean space. Let *M* be a topological *m*-manifold and *N* be a topological *n*-manifold, then *M* × *N* (Cartesian product of *M* and *N*) is a topological (*m* + *n*)-manifold.

This lemma explains the blessing of dimensionality in that the manifold structure is easier to discover in a high-dimensional space. Note that manifold learning in a higher-dimensional space requires more training data. For example, the Cartesian product of a horizontal edge and a vertical edge will produce several combinations including “T”, “+”, “⊢”, “⊣”, and “⊥”. Through combination coding, our direct-fit model stores different patterns like k-d tree (i.e., the centroids of vector quantization or dictionary atoms of sparse coding) but the combination of those patterns will be further enumerated through product topology to support the direct-fit at the higher hierarchy. This enumeration perspective differs from our approach from the HMAX model [81] in which no discrimination was considered for the combination of basic patterns. We argue that the combinatorial coding argument is consistent with recently developed direct-fit model [34], as we will elaborate next.

#### C. Hierarchical Direct-Fit Model with Manifold-based Novelty Detection

**Hierarchical Model Construction** combines layers of simple and complex cells such as HMAX [81] or neocognitron [31], but with an important distinction. The network architecture of our model is not convergent, but *divergent* - one way of generalizing is to still use pooling layers, but we will consider many parallel pooling layers simultaneously. This divergent architecture is inspired by the hypothesis of redundancy exploitation [9] advocated by H. Barlow, since the number of neurons does not decrease, but increases significantly as we move to the higher level of the neocortex. It is natural to ask why higher functions in visual recognition need more neurons. In addition to the argument with combination coding, we note that achieving spatial invariance by max-pooling loses the resolution. To compensate for this sacrifice, context aggregation by dilated convolutions [97] has been developed for semantic segmentation. Mathematically, a dilated convolution is equivalent to a convolution without the follow-up max-pooling operation. Alternatively, we can still use max-pooling, but consider the generalization of dilated convolutions to deformable convolutions [21]. To achieve invariance to local geometric transformations, we can consider a pool of deformable models instead of a single one. Such a combination of space composition with deformable kernels allows us to generate invariant representations with product topology in a bottom-up fashion.

Based on the constructed hierarchical model, we note that the combination of space partitioning (by simple cells) and space composition (by complex cells) can be recursively applied to obtain an over-parameterized model in higher-dimensional space. This recursion is conceptually similar to multi-resolution analysis [57] but operates in a data-adaptive manner (note that we have given up the structures of basis function and inner product in the Hilbert space). At each level, the concatenation of simple and complex cells will map the visual stimuli onto a sequence of invariant representations with increasing dimensionality (field of view). Due to the product-manifold lemma, the number of training data required by our direct-fit model will not exponentially increase (thus, avoiding the curse of dimensionality). Instead, a hierarchical model allows us to automatically adapt to low-dimensional structures in high-dimensional space by extending the k-d tree as follows.

### Product Manifold Tree/Forest

The product manifold tree (pm tree) can be defined as the dual operation of the classic k-d/rp tree. Instead of space partitioning, we recursively merge low-dimensional manifolds in subspaces into higher-dimensional manifolds through product topology. A pm forest uses a pm tree as its building block and consists of an ensemble of pm trees.

How to directly fit the data to the pm tree? Such a problem has been formulated as unsupervised learning in the literature of ML [41, 10]. Unfortunately, all existing work on unsupervised learning and clustering analysis assumes a fixed dimensionality with the input data and focuses on the learning of a distance metric. Our construction of the pm tree attempts to overcome such a barrier by generalizing k-d tree-based clustering with product topology. The performance of nearest-neighbor (NN)-search is known to degrade in high-dimensional space partially due to the surprising behavior of distance metric as dimensionality increases [4]. However, such limitation can be overcome by using the following lemma (note that we do not assume any definition of a distance metric in a topological manifold).

#### Lemma 3

Unsupervised Clustering Lemma on PM Tree

Let **Z** = **X** × **Y** denote the Cartesian product of two topological manifolds. For a vector *z* = [*xy*], its neighborhood search can be carried out by taking the intersection of neighborhoods of *x* ∈ **X** and *y* ∈ **Y**, respectively.

**Sketch of the Proof**. It is known that the subspaces and products of the Hausdorff spaces are Hausdorff. One property of product topology is that if ℬ_*i*_ is a basis for the topology of **X**_*i*_, then the set {**B**_1_ × … × **B**_*n*_ : **B**_*i*_ ∈ ℬ_*i*_} is a basis for the product topology on **X**_1_ × … × **X**_*n*_ [54]. The result follows directly from the definition of a neighborhood basis in the topological manifold.

Such hierarchical construction with interlaced simple and complex cell layers allows an organism to directly fit the visual stimuli into the hierarchical model in a bottom-up fashion. However, such a feedforward process alone is insufficient; it requires a control mechanism (negative feedback) for stability and an escape mechanism (novelty detection) for adaptation. The hippocampus plays an important role in memory consolidation (both old and new) and novelty detection [49]. One way of abstracting the functionality of the hippocampus in visual perception is that it serves the dual role: 1) a control mechanism for implementing negative feedback control loops; 2) an escape mechanism for out-of-distribution (OOD) samples. The first role has been extensively studied in the literature on deep learning (for example, the backpropagation algorithm [83]). Computational modeling of novelty detection includes statistical approaches [59] and neural network-based approaches [60]. Under the framework of our direct-fit hierarchical model, we can revisit these two mechanisms as follows.

### Manifold-based Novelty Detection

In previous work [77], novelty detection was formulated as a twist on the manifold learning problem. First, it linearizes the parameterized manifold, capturing the underlying structure of the inlier distribution. The novelty score is then obtained by factoring in the probability and calculated with respect to the local coordinates of the tangent space of the manifold. However, the task of manifold learning is undertaken by an adversarial auto-encoder, which makes strong assumptions about the input images (e.g., fixed dimensions and inlier/outlier categories). By replacing the adversarial autoencoder with our newly constructed hierarchical model, we can achieve an improved generalization property in the following aspects. First, our construction of a product-manifold tree belongs to unsupervised learning because it directly fits the input data to a pre-constructed data structure. Second, similar to the k-d tree but working toward the opposite direction (increasing dimensionality), the product manifold tree can dynamically accommodate novel events through continual learning. The escape mechanism allows us to incorporate a novel object into the tree so that it is treated as normal in the future.

## IV. Sparse Coding Interpretation: DP-like Solution to 𝓁_∞_-Optimization and Approximate Nearest Neighbor Search

### A. Hierarchical Convolutional Sparse Coding

It is well known that sparse coding offers a powerful analysis of the mechanism of V1 [70, 71]. Meanwhile, sparse representations have also found promising applications in unsupervised learning, such as K-SVD dictionary learning [5] and non-negative matrix factorization [53]. Unlike predetermined dictionaries (e.g., wavelet [57]), data-adaptive dictionary learning such as K-SVD based sparseland [5] has led to a multi-layer formulation of convolutional sparse coding (ML-CSC) [73, 74], which provided an attractive new theoretical framework for analyzing CNN.

#### Multi-Layer Convolutional Sparse Coding (ML-CSC)

The new insight brought about by ML-CSC [73, 74] is to generalize the original sparse coding in a hierarchical (multi-layer) manner. Specifically, a multilayer convolutional sparse coding (ML-CSC) model can be constructed as follows. Suppose that **X** is the input signal and a set of dictionaries is given by 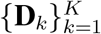 where **D**_*k*_ denotes the dictionary at the level *k*. Then an ML-CSC model can be written as: **X** = **D**_1_ **г**_1_, **г**_1_ = **D**_2_ **г**_2_, …, **г**_*K*−1_ = **D**_*K*_ **г**_*K*_ where **г**_*i*_ = [***w***_1_, …, ***w***_*k*_] denotes the sparse coefficients at the level *i*. Following the convex approximation of 𝓁_0_-optimization in [73], a layered thresholding algorithm runs recursively as follows.

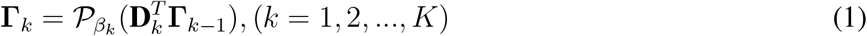

where 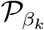 is the standard thresholding operator and 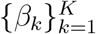 is the set of thresholds at level *k*. As shown in [74], the ML-CSC model manages to decompose a signal **X** ∈ *R*^*N*^ into the superposition of multiple dictionaries **D** = **D**_1_**D**_2_…**D**_*K*_ but the concatenation of these atoms, although it is overcomplete, remains in the space of the same dimension *R*^*N*^. Unfortunately, both K-SVD and ML-CSC are still constructed within the Hilbert space without a change of dimensionality. To the best of our knowledge, no previous work exists in the open literature that extends data-adaptive sparse representation into varying dimensions. Note that wavelet theory achieves the objective of multi-resolution analysis by varying the support of basis functions. Our intuition is that a similar objective can be achieved for sparse data-adaptive representations if we can automatically adapt dictionary learning to the intrinsic low-dimensional structure of the data, such as rp trees [22]. A different way of generalizing the CSC is to construct a dictionary hierarchy with varying dimensionality.

Parallel to the product-manifold lemma, we are interested in developing a recursive strategy to decompose a high-dimensional sparse coding problem into the “product” of multiple lower-dimensional ones. Note that unlike existing work on wavelet decomposition dealing with basis construction in a Hilbert space, we opt to work with directly fitting the data, which is closer to the sparse PCA [99] and K-SVD [5]. Instead of formulating the dictionary learning in the Hilbert space with a fixed dimension (i.e., only vary dictionary size), we imagine a dynamic programming (DP)-like approach to dictionary learning from optimal substructures. The theoretical foundation of such a DP-like solution is the product manifold lemma; and sparse coding offers a well-established framework for exploiting the manifold constraint. We start from the following construction.

#### Hierarchical Convolutional Sparse Coding

Let **X** = **D**_*x*_ **г**_*x*_ and **Y** = **D**_*y*_ **г**_*y*_ denote two dictionary coding schemes wilth **D**_*x*_, **D**_*y*_ ∈ *R*^*n*×*m*^, (*m > n*). Then we start with a coding scheme in the direct-sum space *R*^2*n*^ by 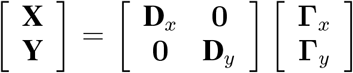 and improve the sparsity by basis pursuit algorithm [19] using the composite dictionary [**D**_*x,i*_ **D**_*y,j*_], (1≤ *i, j* ≤ *m*) (total *m*^2^ atoms). Such process of basis composition can be recursively applied to obtain sparse bases in higher-dimensional spaces (i.e., *R*^*n*^ → *R*^2*n*^ → *R*^4*n*^…).

Our hierarchical extension of the CSC shares a spirit similar to that of dynamic programming in that optimal substructures (atoms) contribute to the (nearly) optimal solution (molecule). Mathematically, 𝓁_0_-optimization is an NP-hard problem; but in biology, evolution does not have foresight and therefore the hierarchical systems generated by evolution might not have a globally optimal structure. What matters more appears to be the nearly decomposability of complex systems [85], which is closely related to the principle of dynamic programming (DP) [13]. Our intuition is that evolution does not need to pursue a globally optimal solution such as NN, but be satisfied with an approximate yet flexible solution so that the organism can adapt to the constantly evolving environment. Based on this observation, we connect the hierarchical CSC with a DP-like recursive solution to approximate NN (ANN) search next.

### B. Anti-sparse coding for Approximate Nearest Neighbor (ANN) Search

Instead of 𝓁_0_-optimization, we conjecture that 𝓁_∞_-optimization (a.k.a., minimax optimization [36]) is a more appropriate framework for analyzing ventral stream processing for the following reasons. First, the strategy employed by V1 [70] follows Barlow’s redundancy reduction principle [11]; however, Barlow has revised this principle to exploit redundancy in [9]. How to exploit the blessing of dimensionality by our hierarchical CSC calls for a fresh look at the definition of sparsity. Second, redundancy has been extensively exploited in information theory for reliable communication [84]. The neocortex faces a similar challenge of robustness to errors (e.g., sensory deprivation and lesions), especially for the high-level layers responsible for important decisions related to behavior. In the literature, it has been shown in [56, 30] that 𝓁_∞_-optimization leads to the so-called Kashin representation (a.k.a. spread representation [30]) where all coefficients are of the same order of magnitude. Such a class of representations is known to robustly withstand errors in their coefficients. Third, the anti-sparse coding scheme based on 𝓁_∞_-optimization is known to facilitate the search for the approximate nearest neighbor (ANN) [46]. Such an interesting connection implies that it is easier to construct a DP-like solution by decomposing the high-dimensional ANN search problem into several subproblems in projected subspaces.

#### Spread Representation

For a given signal *x* in n-dimensional space with **D**_*x*_ ∈ *R*^*n*×*m*^, (*m* > *n*) representing the dictionary, we consider the following 𝓁_∞_-optimization problem.

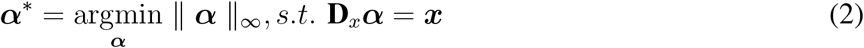

where ‖ ***α*** *‖*_∞_= *max*|*α*_*i*_|, *i* ∈ {1, …, *m*}. The solution to the above 𝓁_∞_-optimization problem [30] boils down to a binarization scheme with components *m* − *n*+1 reaching limits ‖***α*** ‖_∞_ and the remaining *n* − 1 between these two extreme values. This spread representation has led to an anti-sparse coding scheme for the ANN search [46].

#### Anti-sparse coding for ANN search

Given an inquiry vector ***y***, we can find its binarization *e*(***y***) = *sign*(***α****/* ‖ ***α*** ‖_∞_) by solving the anti-sparse coding problem.

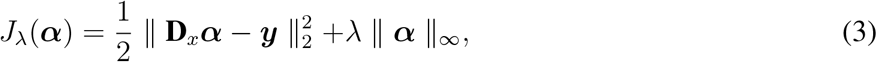

where *λ* is the Lagrangian multiplier. Then the problem of finding an ANN inquiry *z* boils down to

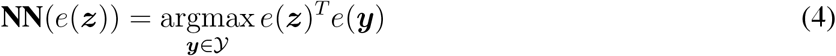

Combining anti-sparse coding with our hierarchical CSC, we can envision a DP-like recursive solution to the ANN search in high-dimensional space. Thanks to product sparse coding lemma, we can recursively construct a dictionary in a high-dimensional space from the direct product of dictionaries in low-dimensional spaces. It follows that an anti-sparse coding scheme in high-dimensional space can be obtained by decomposing it into subproblems in low-dimensional spaces (conceptually similar to the principle of pf dynamic programming). It should be noted that space partitioning, such as the k-d tree and modified checkerboard illusion random projection [95] also supports the NN/ANN search [29]. Therefore, the strategy of ANN search might serve as a common currency to unify top-down and bottom-up processing mechanisms in visual perception.

### C. Theoretical Analysis

#### Relative Invariance to Transformation and Illumination Variation

Unlike wavelet theory [57] in which scale/shift invariance is a property of basis functions, our direct-fit model achieves *relative* invariance to geometric transformations and changes in illumination by learning from the samples. Note that translational invariance is relatively easy to achieve with a maximum-pooling operator with an increased field of view. The invariance in the composition of geometric transformations is beyond the reach of conventional CNN. Our hierarchical model achieves such invariance through densely sampling the data space (guided by ego-motion). In other words, invariant representation arises from the interaction between sensory and motor system - as JJ Gibson comments on visual perception [32], “we see because we move; we move because we see.” Note that such invariance is relative - for rotation that is beyond a critical threshold, the performance of visual recognition degrades rapidly (escape mechanism will be activated).

#### Robustness to Occlusion and Corruption

The robustness of sparse coding recognition [94] is mainly attributed to the use of the 𝓁_1_ norm in convex optimization. The hierarchical extension further improves the robustness by exploiting the redundancy of the representation. As shown in [25], when an overcomplete system is *incoherent*, the optimally sparse approximation to noisy and noiseless data differs from at most a constant multiple of the noise level. Our intuition is that incoherence becomes easier to satisfy in a higher-dimensional space due to the phenomenon of concentration of measures [52]. In addition to the stabilibility results proven in [73], additional robustness to missing and noisy observations comes from the 𝓁_∞_-optimization as rigorously demonstrated by the uncertainty principle in [56].

#### Multiple Stable States

Our new minimax optimization perspective offers a refreshing interpretation about the bistability of anti-sparse coding. Note that ***x*** and −***x*** are always legitimate solutions to the same minimax optimization. This seems a salient difference from 𝓁_*p*_-norm in that 𝓁_∞_-norm is the only one satisfying the symmetry constraint.

## V. Applications to Visual Perception

### A. Grandmother Cell Hypothesis and Face Identity Encoding

There is a long history of debate between localist and distributed representations in psychology and neuroscience [72]. In localist coding, a neuron codes for one familiar thing and does not directly contribute to the representation of anything else [15]. In distributed representations, knowledge is coded in a distributed manner in the mind and brain. The discovery of grandmother cells (a.k.a. Jennifer Aniston cells) [79] can be interpreted as an extreme case of sparseness. The hypothesis of a grandmother cell is also closely related to the binding problem [91] that deals with the integration of individual features into a holistic experience. Our model is in support of the hypothesis regarding the grandmother cell. Following our analysis of the ML-CSC model, the sparsity is expected to increase as the depth of the network increases. This matches our intuition that more abstraction is obtained at higher levels, implying fewer nonzero coefficients required for a global sparse representation. The extreme case will be a single coefficient or neuron. However, it should be noted that there might be massive redundancy in coding any item, such as the grandmother’s face. The locations of these redundant single-neuron encodings can be distributed in the cortex.

Meanwhile, we note that familiarity is an important constraint for the grandmother cell hypothesis. Familiarity and identity of faces are two related but different concepts encoded by the medial temporal lobe (MTL) and the fusiform face area [47]. Face familiarity is related to the bias of visual stimuli; within the framework of sparse coding, faces within the social group (e.g., grandmother’s face) represent the redundancy of training data. This redundancy corresponds to a denser sampling of a local region in the face space, which implies improved sparsity. Therefore, it is easier to observe the firing of single neurons in the class of familiar faces. On the contrary, unfamiliar faces (for example, the cross-race effect [96]) are often more difficult to recognize. This bias is associated with a degraded sparsity due to a poor sampling density. Therefore, it is more difficult to record the response of individual neurons, which is consistent with a recent finding on facial identity coding [17].

### B. Optical Illusion and Context Adaptation

Optical illusions have been widely used by Gestalt psychologists as an experimental tool for studying perceptual organization in visual perception. For example, consider the famous Kanisza’s illusion - an example of an illusory contour [90]. What has been less studied is the perturbed version of this illusion - i.e., as one increases the size of the black triangle and the distance among three pacmacs, it will become more and more difficult to perceive the illusory “white triangle” at the center. Such experimental findings cannot be explained by Gestalt theory because there is no prediction of the critical boundary condition for perceptual organization to fall apart. Our hierarchical direct-fit model can predict that such a threshold is determined by the size of the fovea centralis. When the distance is above this threshold, there is no previous experience (training data) to support the perceptual grouping of white triangles.

Another celebrated illusion related to context adaptation is Ted Adelson’s checkerboard illusion [2] (see Fig. 2b). Two blocks with identical color values can appear to have opposite perceived luminance (brightness). Such an inconsistency is the consequence of perceptual organization; namely, the global constraint of perceiving a checkerboard dictates the interpretation at lower levels. By contrast, if two additional vertical strips having identical color are added, they interfere with the result of perceptual organization at the global level. Consequently, two blocks of the same color as two strips tend to be grouped together.

**Fig. 2:**
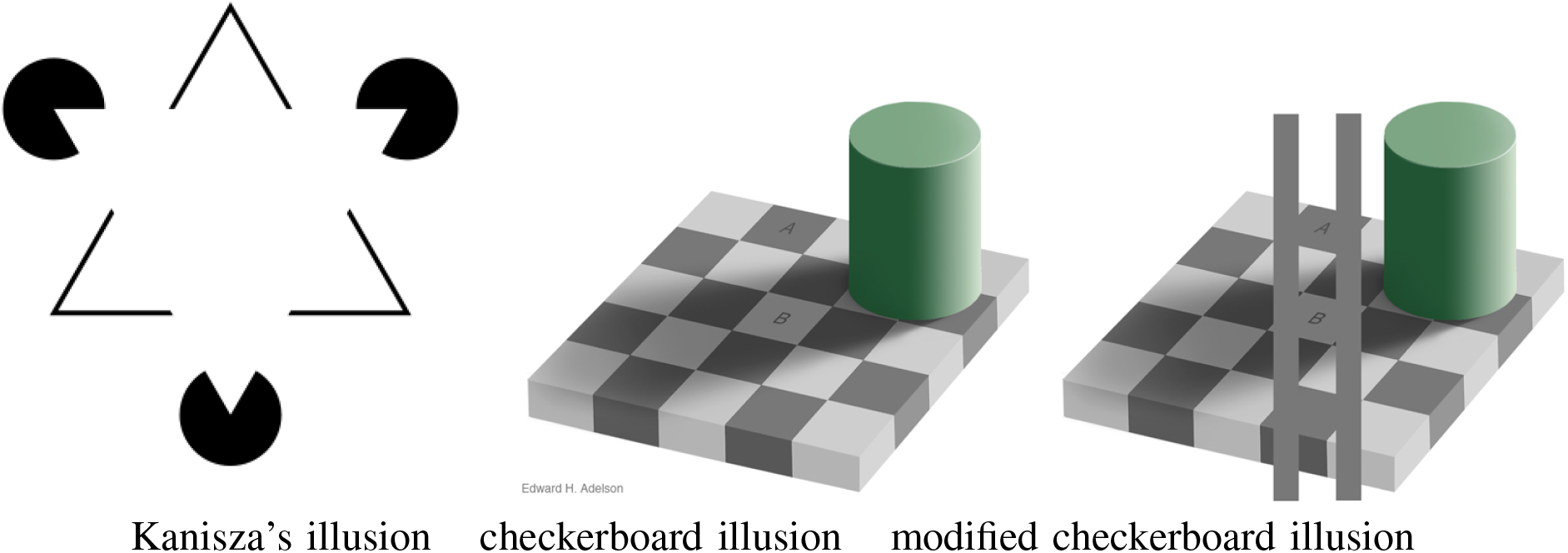
Optical illusion of an illusory contour. The white triangle is perceived as an illusory contour (the illusion will disappear as the size of the triangle increases).

### C. Multi-stable Perception and Novelty Detection

Multistable perception refers to the spontaneous alternation between two (or more) perceptual states when visual stimuli are inherently ambiguous (refer to Fig. 3). Multistable perception is a widely studied topic in visual perception [55] due to its close relationship to sensory awareness or consciousness. The neural basis of multi-stable perception [86] has emphasized the role of high-level brain mechanisms that are involved in actively selecting and interpreting sensory information from lower-level processes. However, no theoretical explanation about the deeper mechanism of multi-stable perception exists, not to mention the prediction about when this phenomenon will occur. Our hierarchical direct-fit model shed some insight to this long-standing open question in that bi-stable perception directly corresponds to *x* and −*x* in our anti-sparse coding formulations. As discussed above, if neocortex is organized in such a way to recursively solve the minimax optimization problem, recursive ANN search from low to high might end up with two equally possible solutions (*x* vs. −*x*).

**Fig. 3:**
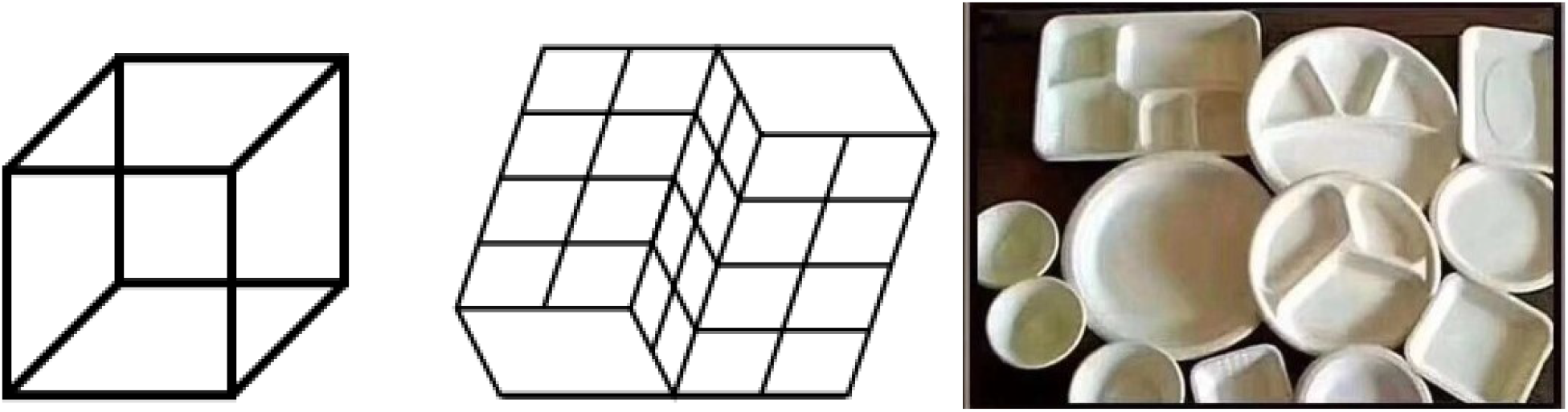
Multi-stable perception (each image admits two competing interpretation results).

Our final experiment deals with novelty detection in visual perception. In the so-called Thatcher effect [1], one can observe that it is relatively easy to detect inverted eyes and mouth from an upright face (see Fig. 4a). This detection of an anomaly (novelty) can be explained away because local patches around the eye and mouth regions are not compatible with the global impression of a human face. However, such novelty detection tasks become much more challenging with the inverted face (see Fig. 4b). Note that an inverted face is a rare event for HVS (e.g., face recognition becomes almost impossible [87]), and the hippocampus can only tell it is a novelty. Accordingly, top-down feedback in a hierarchical model will not detect the inconsistency between the inverted face and the eyes/mouth in normal position.

**Fig. 4:**
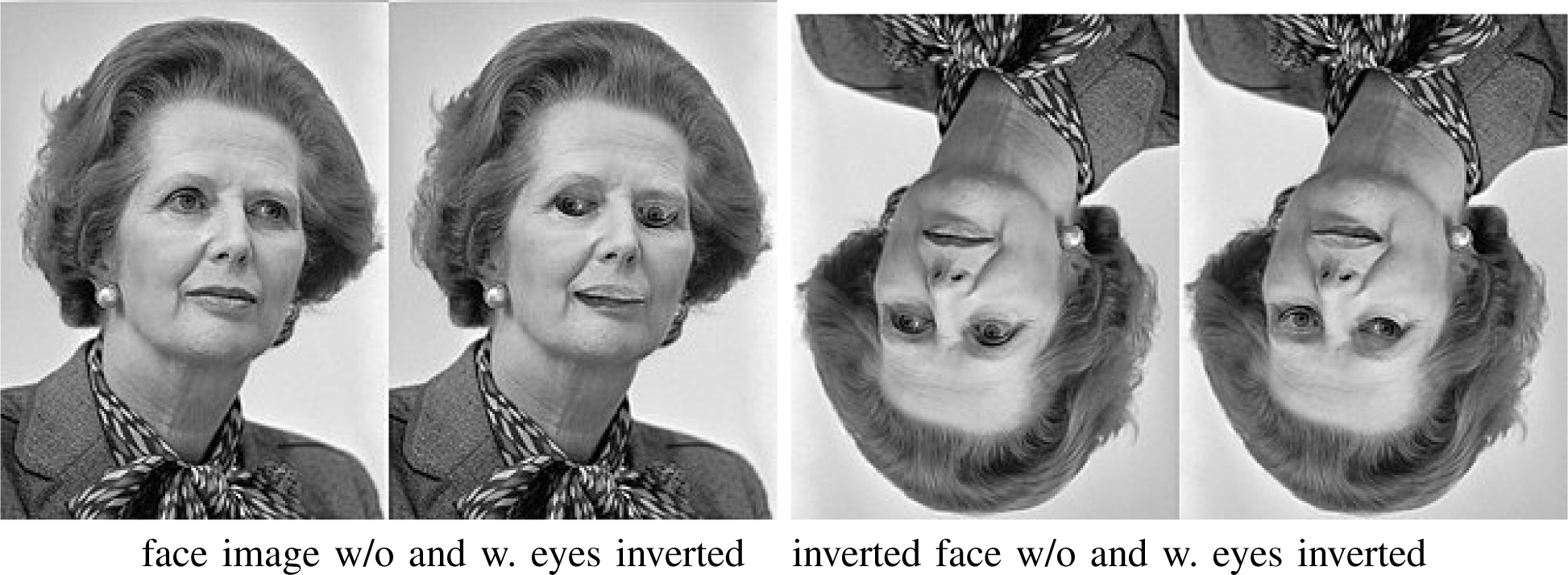
Thatcher effect. It is much easier to detect inverted eyes from an upright face than from an upside-down face.

## VI. Two Possible Implementations Using Artificial Neural Networks

### A. Asymmetrical Autoencoder: Decoder Matters More

The autoencoder represents a popular network architecture for unsupervised learning. A straightforward application of the sparse coding principle to the autoencoder is possible [69]. The hierarchical extension of the CSC inspires us to consider the design of an asymmetrical autoencoder, as shown in Fig. 5. Its asymmetrical design is also biologically inspired by H. Barlow’s redundancy exploitation hypothesis [9]. The hierarchical organization of the neocortex is believed to reflect the nested structure of the physical world [35], indicating that the decoder (responsible for reconstruction of internal representations) plays a more important role than the encoder. More specifically, we can implement a prototype by combining an over-parameterized autoencoder [80] with manifold-based novelty detection [77]. In the literature, autoencoder-based representation learning [88] is known to be capable of learning disentangled and hierarchical representations. Instead of storing training samples as attractors [80], we envision that a hierarchy of increasingly abstract concepts can become attractors of an over-parameterized asymmetrical autoencoder. In the presence of a new category (novelty), the escape mechanism will be activated and the decoder will be further expanded to accommodate the novelty class (i.e., consolidation of new memory).

**Fig. 5:**
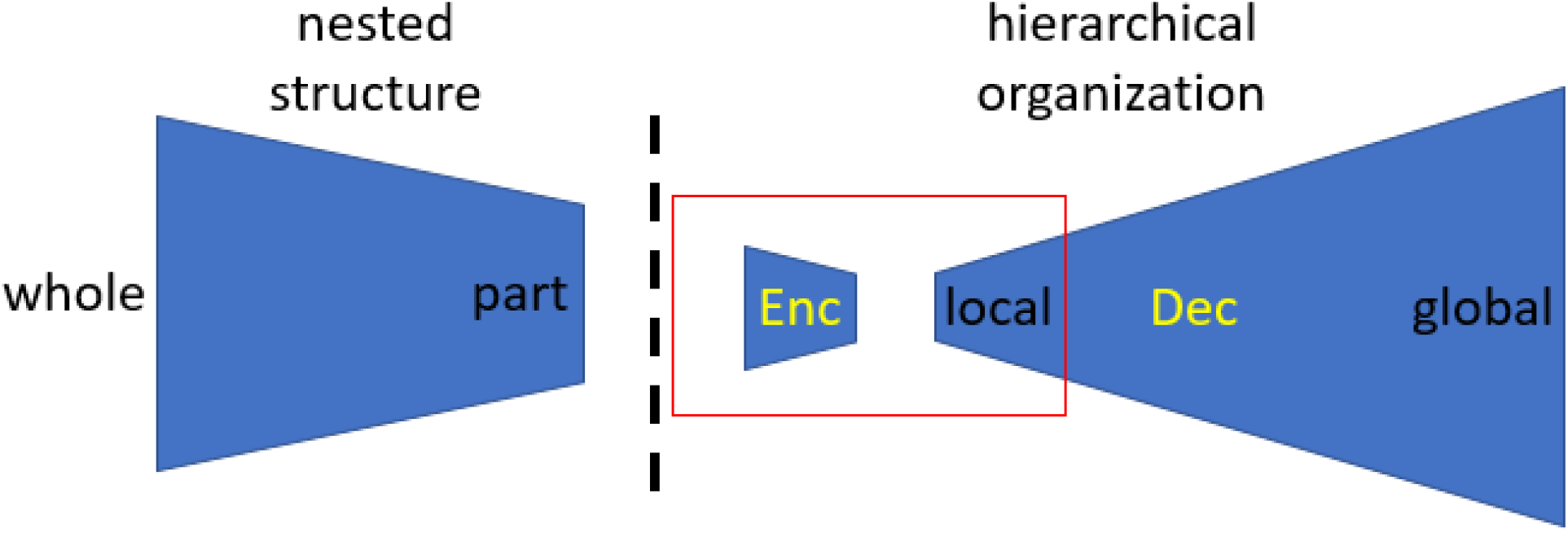
Architecture of an asymmetrical autoencoder implementing the principle that nested structure in the physical world is reflected by the hierarchical organization of the neocortex (the dashed line marks the boundary between environment and organism; red box corresponds to the standard symmetric autoencoder).

Such unsupervised and continual learning can lead to monotonically increased memory capacity for associative memory implemented by overparameterized autoencoder. Unlike [80] treating sequence encoding as composition maps and limit cycles, we argue that a biologically more plausible mechanisms for memory storage is based on the rich interaction between sensory and motor systems. From the direct-fit perspective, motion dictates the regime within which the organism achieves the invariance to the composition of geometric transformations. The rich interaction between sensory and motor systems contributes to the formulation of reconstruction problems at multiple scales from parts to the whole [40] as an unsupervised mechanism of learning invariant representations. When the sensory motion goes out of the normal range (e.g., rotating a book continuously), the asymmetrical autoencoder will fail to recognize the object (i.e., it will be treated as a novelty).

### B. Polychronous Neuronal Group (PNG) and Spiking Neural Networks

In [45], polychronization was conceived as a basic mechanism for computing with spikes. It is built upon Hebb’s postulate, but extends it by relaxing the synchronous firing into polychronous time-locked patterns. Therefore, the group of neurons that are spontaneously organized by the fundamental process of spike-timing-dependent plasticity (STDP) is called Polychronous Neuronal Group (PNG). The mechanism of polychronization has recently been studied in [26, 44] as a plausible solution to the problem of feature binding. Using a spiking neural network (SNN), input training images (sensory stimuli) can be mapped to a hierarchy of PNGs by the emergence of polychronization. A mathematical abstraction of PNG is that it maps some partitioned space to a binary output (firing or no firing).

Unlike previous studies [26, 44], our over-parameterized direct fit model can be implemented on SNN with a divergent architecture, as shown in Fig. 6. Such a divergent architecture directly matches our intuition of abstracting complex cells by space composition. It is easy to see that the number of PNGs can grow exponentially as the number of neurons increases. Why do we need more PNGs at the higher level of the neocortex? The combinational coding argument (the end of Sect. III-B) suggests that PNGs could represent a biological plausible storage mechanism. They are physical implementations of attractors in artificial neural networks. One way to experiment with our divergent SNN is to focus on its memory capacity; the other promising direction is to build a dorsal-ventral SNN to study the binding mechanism between “what” and “where” according to recent work of SNN for object vision [37] and spatial vision [75], respectively.

**Fig. 6:**
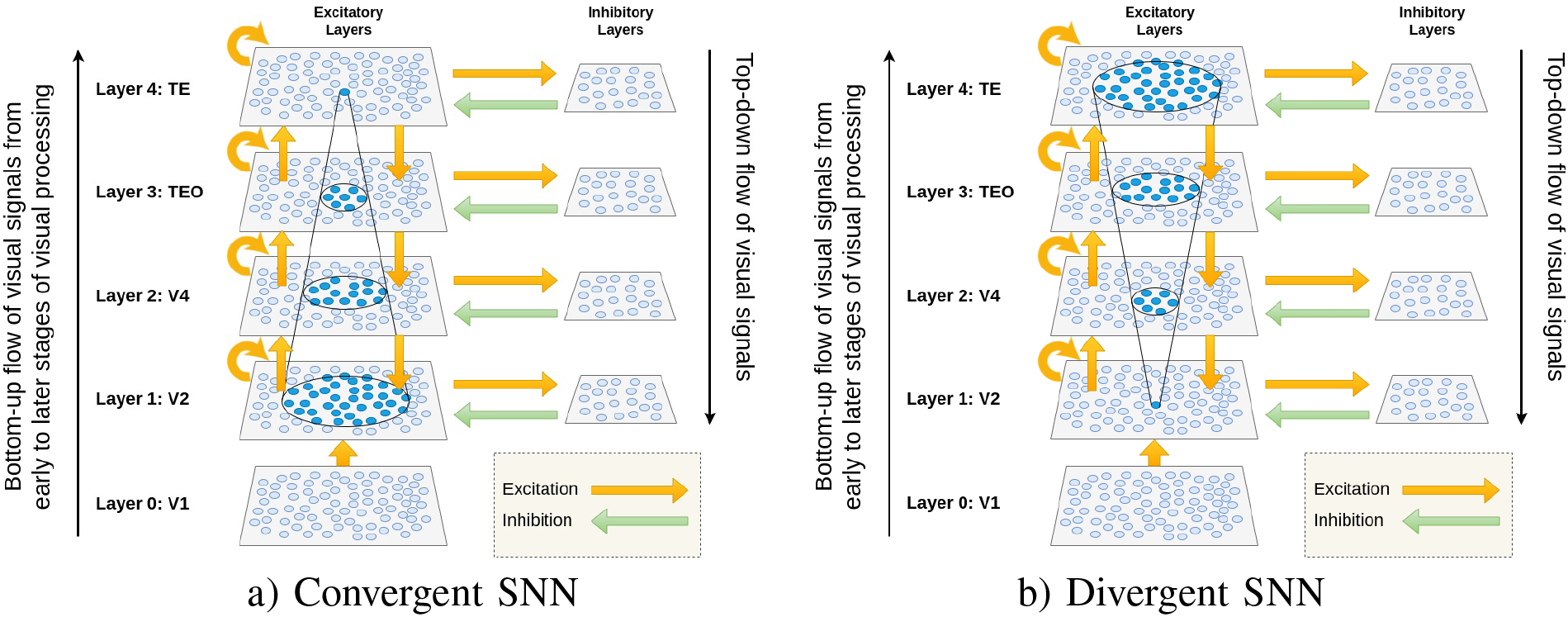
Current implementation of SNN [26, 44] vs. the proposed implementation (there are more cells and synapses at higher levels than at lower levels).

## VII. Conclusions

We challenge the conventional way of modeling simple/complex cells by constructing space partitioning/composition in the topological space. Without imposing additional structures such as distance metric, we can construct product-manifold trees as a novel data structure suitable for the task of direct-fit visual perception. We demonstrate the biological plausibility of our construction and offer a sparse coding interpretation. It is possible to test the developed theory by constructing an asymmetrical autoencoder or a divergent SNN. For asymmetrical autoencoder, it will be interesting to study how a divergent architecture can maximize the capacity of associative memory. For divergent SNN, the objective is to experimentally verify the parallel and sequential binding mechanisms and their associated combination coding arguments, which might offer supporting evidence for the grandmother cell hypothesis.

